# A quantitative pipeline to assess secretion of human leptin coding variants reveals mechanisms underlying leptin deficiencies

**DOI:** 10.1101/2024.03.01.582913

**Authors:** Harry J. M. Baird, Amber S. Shun-Shion, Edson Mendes de Oliveira, Danièle Stalder, Jessica Eden, Joseph E. Chambers, I. Sadaf Farooqi, David C. Gershlick, Daniel J. Fazakerley

## Abstract

The hormone leptin, primarily secreted by adipocytes, plays a crucial role in regulating whole-body energy homeostasis. Homozygous loss-of-function mutations in the leptin gene *(LEP)* cause hyperphagia and severe obesity, primarily through alterations in leptin’s affinity for its receptor or changes in serum leptin concentrations. Although serum concentrations are influenced by various factors (e.g., gene expression, protein synthesis, stability in the serum), proper delivery of leptin from its site of synthesis in the endoplasmic reticulum via the secretory pathway to the extracellular serum is a critical step. However, the regulatory mechanisms and specific machinery involved in this trafficking route, particularly in the context of human *LEP* mutations, remain largely unexplored.

We have employed the Retention Using Selective Hooks (RUSH) system to elucidate the secretory pathway of leptin. We have refined this system into a medium-throughput assay for examining the pathophysiology of a range of obesity-associated *LEP* variants. Our results reveal that leptin follows the default secretory pathway, with no additional regulatory steps identified prior to secretion.

Through screening of leptin variants, we identified three mutations that lead to proteasomal degradation of leptin and one mutant that significantly decreased leptin secretion, likely through aberrant disulfide bond formation. These observations have identified novel pathogenic effects of leptin variants, which can be informative for therapeutics and diagnostics. Finally, our novel quantitative screening platform can be adapted for other secreted proteins.

## Introduction

Obesity is highly heritable (1). The first gene to be associated with severe early-onset obesity in humans was *LEP* (*2*), which encodes leptin, an adipose tissue-derived hormone that regulates whole-body energy homeostasis (3, 4). Congenital leptin deficiency in mouse models (*ob/ob)* and in humans (3, 4) is characterised by undetectable serum leptin and early-onset severe obesity associated with intense hyperphagia (2, 4). In most people, the amount of circulating leptin is influenced by nutrient availability (5) and is proportional to body fat mass (6). Increased adiposity augments circulating leptin through increased *LEP* transcription (7), translation (8–11), and secretion from adipose tissue. However, very little is known about the secretory route of leptin or the effects of human *LEP* variants on leptin secretion.

Leptin loss-of-function missense variants have been described in humans with severe obesity (2, 12–14). These typically result in low serum leptin (2, 12, 15). Another class of leptin deficiency is associated with normal-to-high serum leptin, but these variants are biologically inactive due to structural alterations that affect receptor binding (13). Recently, variants that antagonise the leptin receptor have also been reported (16). In the case of variants resulting in low serum leptin, there are a number of possible mechanisms, including defects in transcription, translation or secretion. Resolving these mechanisms has been challenging due to the technical difficulty of interrogating secretory kinetics.

Here, we have developed a platform to overcome these challenges and interrogate how natural coding variants of *LEP* affect circulating leptin. We developed a Retention Using Selective Hooks (RUSH)-based 96-well plate assay where we trap newly synthesised leptin in the endoplasmic reticulum (ER) and monitor secretion from cells following its synchronised release (17). Using this assay, we found that leptin is secreted via the classical ER-Golgi-plasma membrane pathway in both HeLa cells and 3T3-L1 adipocytes, and we screened naturally occurring leptin variants for effects on leptin expression and/or secretion. Four variants of interest lowered leptin secretion; three caused leptin degradation, and one specifically slowed the kinetics of leptin export from the ER.

## Results

### Establishing cell models to study leptin secretion

Leptin is a soluble protein constitutively secreted from adipocytes (18). Constitutively secreted proteins are synthesised in the ER, trafficked to the Golgi apparatus, and packaged into secretory carriers which fuse with the cell surface (19). As secreted proteins are continuously lost from the cell, uncoupling secretion kinetics from transcription and translation is challenging. To overcome this, we have developed a quantitative kinetic assay for leptin secretion using the Retention Using Selective Hooks (RUSH) system (17, 19). RUSH allows retention of a protein-of-interest within the ER after synthesis and its synchronous release through the secretory pathway upon addition of biotin. To study the secretion of leptin using RUSH, we generated a stable HeLa cell line expressing a synthetic *LEP* bicistronic construct to produce a leptin fusion protein with SBP and HaloTag (SBP-HaloTag-leptin) (**Fig. 1A**), and streptavidin-KDEL. The SBP-HaloTag-leptin fusion is retained in the ER and can be visualised using fluorescence microscopy through labelling with HaloTag-JFX647 dye. The addition of biotin releases SBP-HaloTag-leptin from the ER along its trafficking pathway before being secreted from the cell into the extracellular media (**Fig. 1**B).

**Figure 1:**
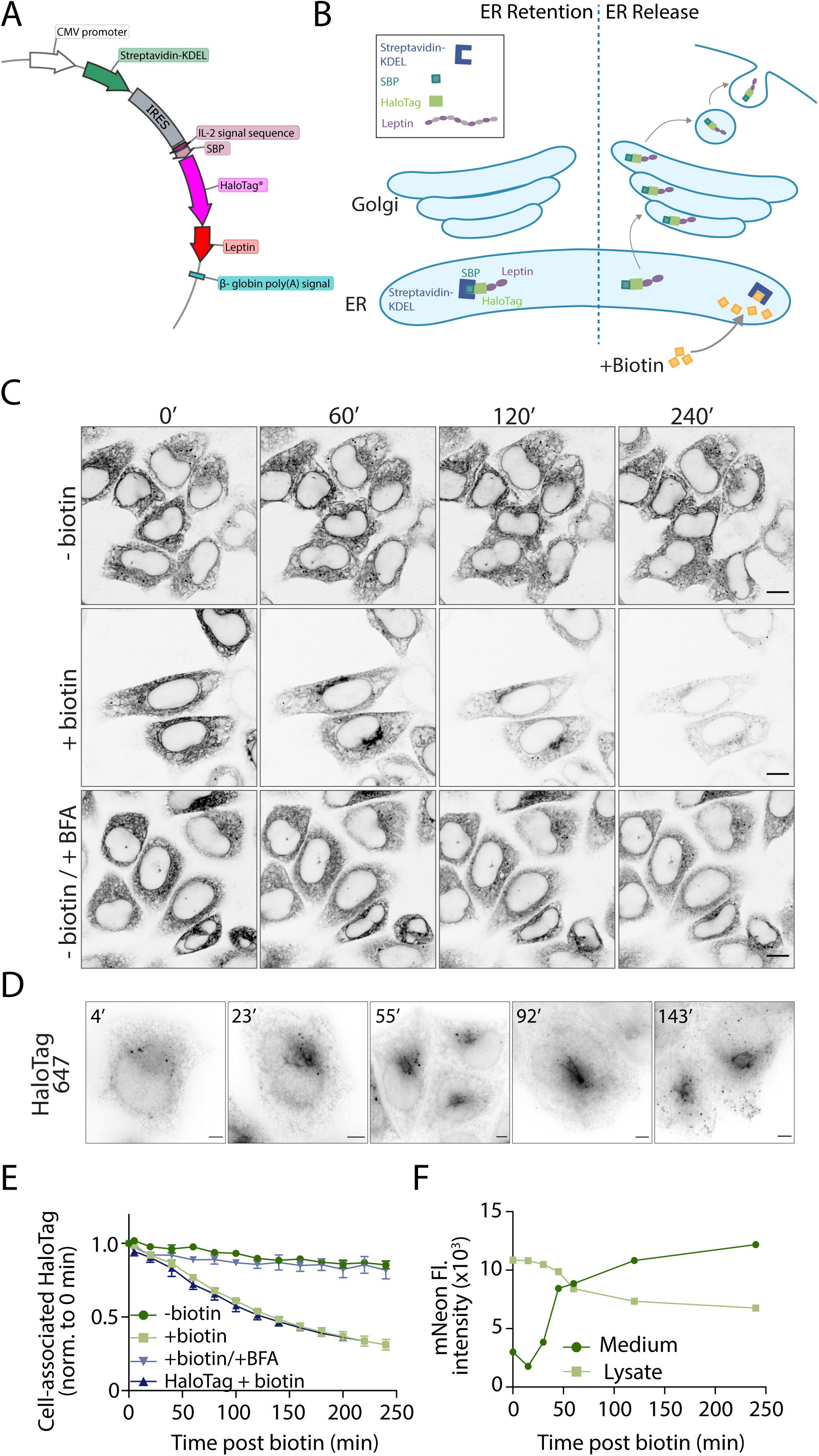
Development and validation of a RUSH-based kinetic secretory assay to study leptin secretion. A) Plasmid map of the piggybac vector used for stable cell line generation. A bicistronic internal ribosome entry site (IRES) vector is driven by the CMV promoter expressing an upstream streptavidin-KDEL and downstream a fusion protein of the signal peptide from IL-2, a HaloTag and leptin. B) Schematic of the RUSH-based system to monitor leptin secretion. Prior to biotin treatment, SBP-HaloTag-leptin is retained in the ER through an interaction between the SBP and the Strep-KDEL. Upon biotin addition, biotin outcompetes the SBP interaction and releases SBP-HaloTag-leptin from the ER, through the Golgi for secretion. C) Live cell imaging of SBP-HaloTag-leptin in HeLa cells labelled with JFX646 Halo dye. Cells were treated with biotin and BFA where indicated and imaged at 0, 60, 120 and 240 min after biotin addition. Representative of n = 2; scale bar: 10 µM. D) Live cell super-resolution imaging of SBP-HaloTag-leptin in HeLa cells labelled with JFX646 Halo dye. Images are from specified time points following biotin addition. Representative of n = 3; scale bar: 5 µm. E) Kinetic analysis of SBP-HaloTag-leptin secretion using live cell imaging. Cells were treated with biotin and/or BFA where indicated. SBP-HaloTag was included as a control in this experiment (ctrl: dark blue). Fluorescence intensity of cell-associated HaloTag signal was measured and normalised to the 0 min time point. n = 3 for SBP-Leptin-HaloTag cells +/- biotin; n = 2 for SBP-Leptin-HaloTag cells + biotin/BFA and SBP-Halotag cells (ctrl). F) SBP-mNeonGreen-leptin HeLa cells were treated with biotin for specified time periods and the mNeonGreen fluorescence intensity in culture media (dark green) and cell lysate (light green) measured. n = 1.

To validate the use of this assay to study leptin secretion, we used live-cell fluorescence confocal imaging to visualise and quantify intracellular leptin over 4 h (**Fig. 1C**). Cells were incubated with HaloTag ligand JFX646 prior to the experiment to label the tagged leptin. Cells were then treated with biotin and imaged at 20 min time intervals for 240 min. Without biotin, SBP-HaloTag-leptin remained in the ER for the duration of the experiment. At 60 min after biotin treatment, SBP-HaloTag-leptin accumulated in the Golgi (**Fig. 1C**). By 120 and 240 min, we observed a decrease in cellular fluorescence, suggesting that the SBP-HaloTag-leptin had been secreted (**Fig. 1C**). These observations were confirmed using live cell structured-illumination microscopy, where we observed that SBP-HaloTag-leptin accumulated in the Golgi 23 min after biotin addition (**Fig. 1D**). After 55-92 min of biotin treatment, SBP-HaloTag-leptin was localised to the Golgi and to small vesicles which appeared closer to the cell periphery after 143 min (**Fig. 1D**).

Analysis of the kinetics of SBP-HaloTag-leptin secretion by measuring the cell-associated HaloTag-JFX646 signal and normalising to HaloTag-JFX646 fluorescence at 0 min revealed 68% loss of intracellular leptin signal over 4 h after biotin addition (**Fig. 1E).** This loss of SBP-HaloTag-leptin signal was abrogated by treatment with the ER-to-Golgi trafficking inhibitor Brefeldin A (BFA) (**Fig. 1C**, quantified in **Fig. 1E**), suggesting that SBP-HaloTag-leptin is secreted via the classical ER-Golgi pathway. Additionally, the secretory kinetics of SBP-HaloTag-leptin were comparable to an ER-lumen localised SBP-HaloTag-only control, suggesting that leptin-SBP-HaloTag secretion occurs constitutively and that signals within leptin do not confer additional regulation (**Fig. 1E**). This imaging, together with the quantitative data, suggests that after release from the ER, SBP-HaloTag-leptin traffics with typical constitutive secretory kinetics from the ER to the Golgi. At the Golgi, it is sorted into small vesicles and subsequently secreted from the cells.

Determining leptin secretory kinetics using the confocal microscopy-based assay described above relies on measuring signal loss from cells as SBP-HaloTag-leptin is secreted. To confirm leptin is secreted from cells rather than degraded intracellularly, we directly measured leptin secreted into the culture medium using cells expressing SBP-mNeonGreen-leptin. Cells were treated with biotin for specific time periods up to 240 min, after which the culture media was collected and the cells lysed. The mNeonGreen fluorescence intensity of the cell culture media and the cell lysates were quantified as a measure of SBP-mNeonGreen-leptin secretion. The fluorescent intensity of the media increased >3-fold from 0 to 240 min, concomitant with a 37% decrease in fluorescent intensity in the cell lysate. These data suggest that SBP-mNeonGreen-leptin is secreted from cells into the cell culture media (**Fig. 1F**). Overall, our findings demonstrate that the RUSH system can be used to study the secretory route and kinetics of leptin and that leptin follows a classical constitutive secretory pathway.

We next established a more physiologically relevant 3T3-L1 adipocyte cell line to study leptin secretion using the same SBP-HaloTag-leptin/Streptavidin-KDEL system. Results using the adipocyte model were consistent with the HeLa model, including the transit through the secretory pathway (**Fig. 2A, B**), the sensitivity to BFA (**Fig. 2A**), and leptin secretion from the cell to the media (**Fig. 2A&C**). Together, these data suggest that we can assess leptin secretion using this RUSH-based system in 3T3-L1 adipocyte cells and that leptin traffics through a conventional constitutive secretory pathway. This is in agreement with earlier observations made in HeLa cells.

**Figure 2:**
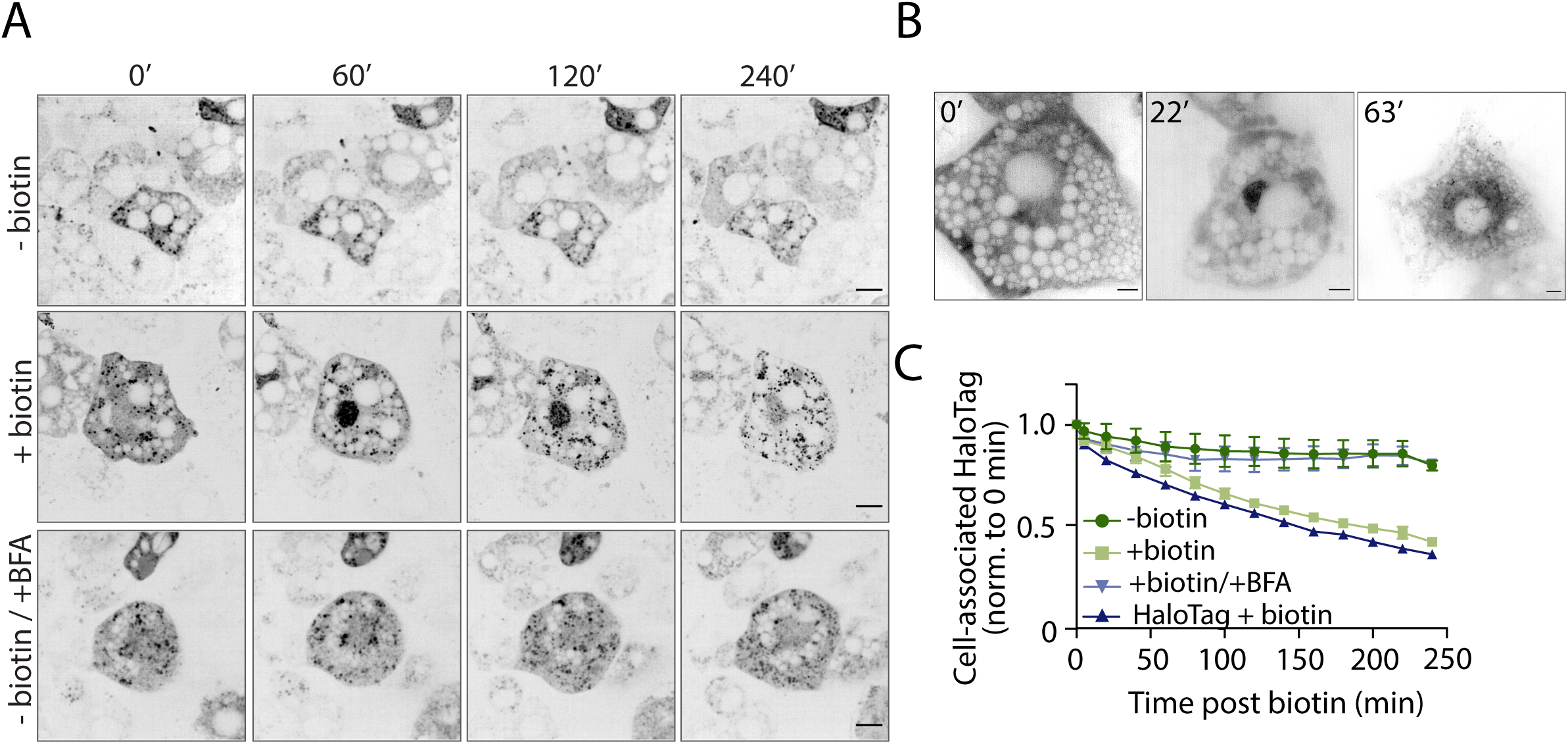
Establishing the RUSH-based assay for leptin secretion in 3T3-L1 adipocytes. A) Live cell imaging of SBP-HaloTag-leptin in 3T3-L1 adipocytes labelled with JFX646 Halo dye. Cells were treated with biotin and/or BFA and imaged at 0, 60, 120 and 240 min. Representative images from n = 3; scale bar: 10 µm. B) Live cell super-resolution imaging of SBP-HaloTag-leptin in 3T3-L1 adipocytes labelled with JFX646 Halo dye. Images are from specified time points following biotin addition. Representative of n = 3; scale bar: 5 µm. C) Kinetic analysis of SBP-HaloTag-leptin secretion using live cell imaging. Cells were treated with biotin and/or BFA where indicated. SBP-HaloTag was included as a control in this experiment (ctrl: dark blue). Fluorescent intensity of cell-associated Halo signal was measured and normalised to the 0 min time point. n = 3 for SBP-Leptin-HaloTag cells +/-biotin +/- BFA; n = 1 for SBP-HaloTag cells (ctrl).

### Screening human LEP coding variants for effects on expression and secretion

Low serum leptin contributes to the development of severe obesity (2, 20). Human coding variants may cause lower serum leptin through decreased *LEP* mRNA abundance or translation, protein degradation, decreased serum stability of the protein product, or impaired leptin secretion from adipose tissue. Despite the importance of leptin to energy homeostasis in humans, there has been relatively little investigation of the mechanisms explaining low serum leptin of *LEP* coding variants. Having established a quantitative system for studying leptin secretion, we conducted a blinded screen of 12 human *LEP* variants (**Fig. 3A**; **Table 1**) to determine whether these were associated with impaired leptin production or secretion. The variants selected were linked to low serum leptin (p.L72S, p.N103K, p.R105W, p.C117Y, p.P23R)(12, 21–24) and/or those associated with obesity with a less clear or debated mechanism-of-action (p.H118L, p.S141C, p.D100N) (25, 26) (**Table 1**). As negative controls, we included three variants (p.G59S, p.P64S, p.D100Y) (13, 16) previously characterised as biologically inactive with no secretory phenotype, and one variant (p.V110M) found in an overweight (BMI 27.2) individual with normal serum leptin (27). Each variant was cloned into a piggyBac RUSH plasmid system (28, 29), and 12 stable cell lines were generated for analysis. Using these cell lines with appropriate wildtype (WT) controls, we measured cell-associated SBP-HaloTag-leptin by live cell imaging over 4 h.

**Figure 3:**
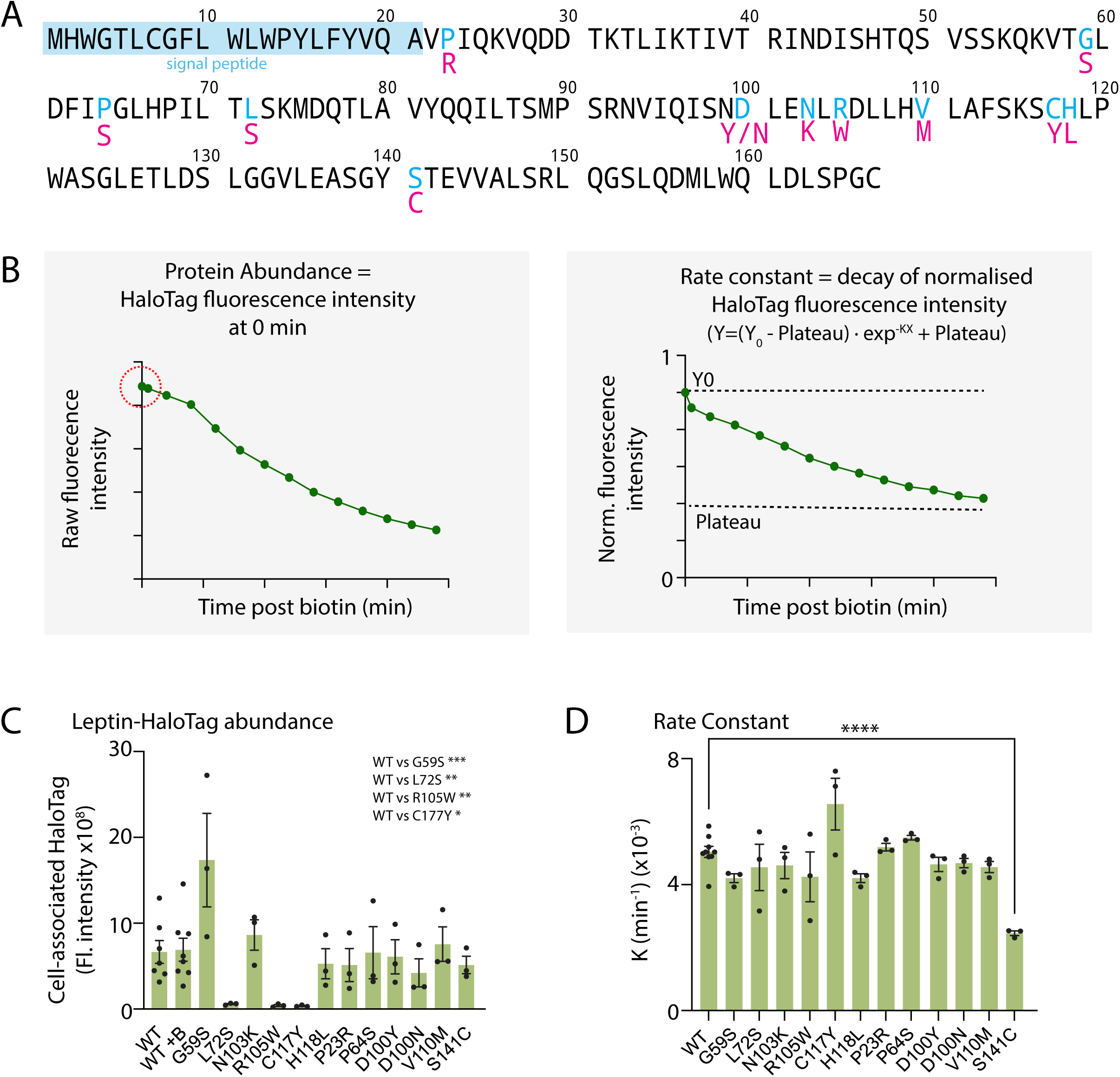
Screening of natural Leptin variants using the Leptin RUSH-based assay. A) Schematic of the human leptin variants used in this screen. Blue residues indicate the site of mutation and red residues underneath indicate the amino acid substitution. B) Schematic of data processing and interpretation. For each mutant, time-course RUSH assays were performed. The first data point collected was the raw initial fluorescent intensity of the construct at time point 0’ as a readout of protein abundance (circled in red). The data were subsequently normalised to 0 min time point and the rate constant K was calculated for SBP-HaloTag-leptin secretory kinetics (depletion of HaloTag signal). C) Raw initial 647 fluorescence values at time point 0’ for all SBP-HaloTag-leptin constructs screened. n= 7 for WT SBP-HaloTag-leptin; n = 3 for all leptin variants. Statistical analysis comparing WT vs each variant performed by 2-way ANOVA. Post-hoc tests were corrected for multiple comparisons (two-sided Dunnett’s test), * *p* value <0.05, ** *p* value <0.01, *** *p* value <0.001. D) Rate constant K values for SBP-HaloTag-leptin variants. n = 7 for WT SBP-HaloTag-leptin; n = 3 for all leptin variants. Statistical analysis comparing WT vs each variant performed by 2-way ANOVA. Post-hoc tests were corrected for multiple comparisons (two-sided Dunnett’s test), **** *p* value <0.0001.

**Table 1:**
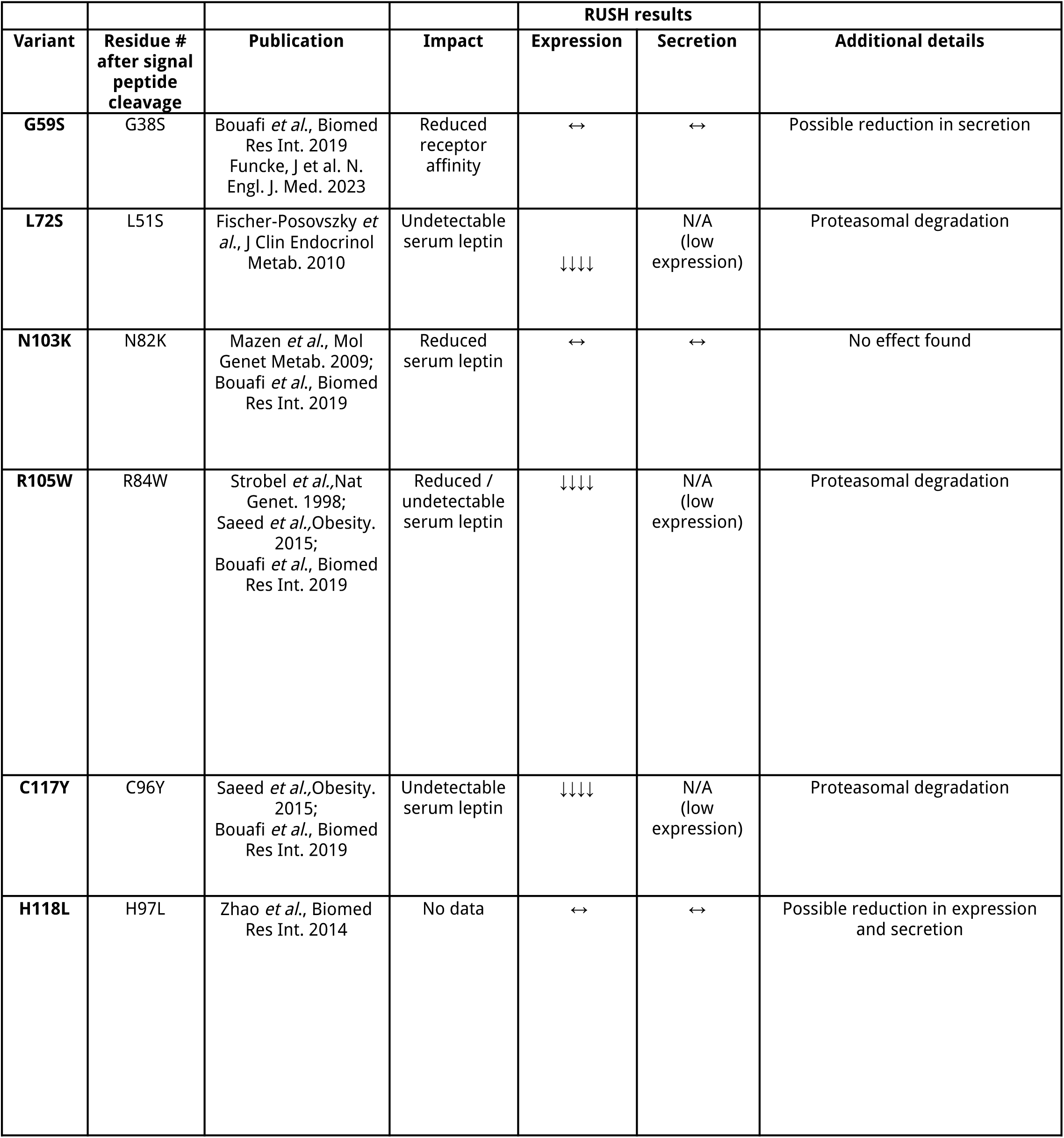
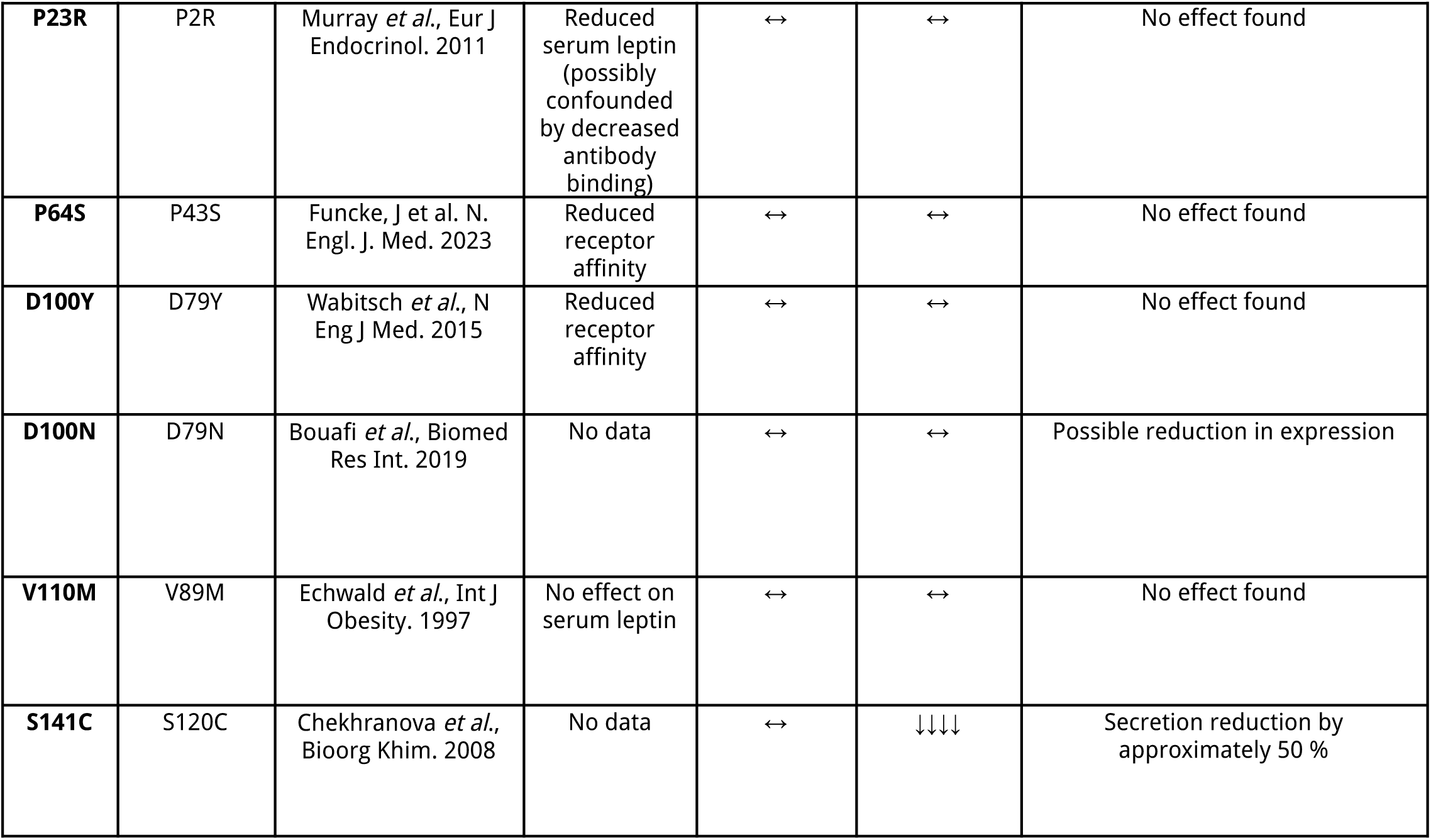
Table of leptin variants studied.

| Variant | Residue #<br>after signal<br>peptide<br>cleavage | Publication | Impact | RUSH results |  | Additional details |
| --- | --- | --- | --- | --- | --- | --- |
|  |  |  |  | Expression | Secretion |  |
| **G59S** | G38S | Bouafi *et al.*, Biomed Res Int. 2019<br>Funcke, J et al. N. Engl. J. Med. 2023 | Reduced receptor affinity | ↔ | ↔ | Possible reduction in secretion |
| **L72S** | L51S | Fischer-Posovszky *et al.*, J Clin Endocrinol Metab. 2010 | Undetectable serum leptin | ↓↓↓↓ | N/A (low expression) | Proteasomal degradation |
| **N103K** | N82K | Mazen *et al.*, Mol Genet Metab. 2009;<br>Bouafi *et al.*, Biomed Res Int. 2019 | Reduced serum leptin | ↔ | ↔ | No effect found |
| **R105W** | R84W | Strobel *et al.*, Nat Genet. 1998;<br>Saeed *et al.*, Obesity. 2015;<br>Bouafi *et al.*, Biomed Res Int. 2019 | Reduced / undetectable serum leptin | ↓↓↓↓ | N/A (low expression) | Proteasomal degradation |
| **C117Y** | C96Y | Saeed *et al.*, Obesity. 2015;<br>Bouafi *et al.*, Biomed Res Int. 2019 | Undetectable serum leptin | ↓↓↓↓ | N/A (low expression) | Proteasomal degradation |
| **H118L** | H97L | Zhao *et al.*, Biomed Res Int. 2014 | No data | ↔ | ↔ | Possible reduction in expression and secretion |
| **P23R** | P2R | Murray *et al.*, Eur J Endocrinol. 2011 | Reduced serum leptin (possibly confounded by decreased antibody binding) | ↔ | ↔ | No effect found |
| **P64S** | P43S | Funcke, J et al. N. Engl. J. Med. 2023 | Reduced receptor affinity | ↔ | ↔ | No effect found |
| **D100Y** | D79Y | Wabitsch *et al.*, N Eng J Med. 2015 | Reduced receptor affinity | ↔ | ↔ | No effect found |
| **D100N** | D79N | Bouafi *et al.*, Biomed Res Int. 2019 | No data | ↔ | ↔ | Possible reduction in expression |
| **V110M** | V89M | Echwald *et al.*, Int J Obesity. 1997 | No effect on serum leptin | ↔ | ↔ | No effect found |
| **S141C** | S120C | Chekhranova *et al.*, Bioorg Khim. 2008 | No data | ↔ | ↓↓↓↓ | Secretion reduction by approximately 50 % |

These imaging data were used to generate two metrics: 1) the raw initial fluorescence intensity value before normalisation, as a measure of variant protein abundance in cells (**Fig. 3B-left panel**); and 2) a rate constant (K) for SBP-HaloTag-leptin secretion (i.e., the decay of fluorescence intensity during the assay) calculated after normalisation of fluorescence intensity data to cell fluorescence at 0 min (**Fig. 3B-right panel**).

Most leptin variants were expressed and secreted with similar kinetics to WT leptin (**Fig. 3C&D; Supplemental Fig. 1**). However, there were four variants-of-interest. p.L72S, p.R105W and p.C117Y substantially reduced initial fluorescent intensity compared to WT leptin (**Fig. 3C**). Interestingly, p.S141C had normal abundance but a reduced secretory rate constant (**Fig. 3D**), suggesting that the p.S141C variant induces a specific secretory defect.

### Specific leptin variants are degraded via the proteasome

Previous data suggests that subjects with the p.L72S, p.R105W, and p.C117Y variants have low or undetectable leptin serum (21, 23). Since the secretory kinetics of these variants were largely unaffected (**Supplemental Fig. 1**), p.L72S, p.R105W, and p.C117Y variants likely lower serum leptin through decreased leptin protein abundance (**Fig. 4A)**.

**Figure 4:**
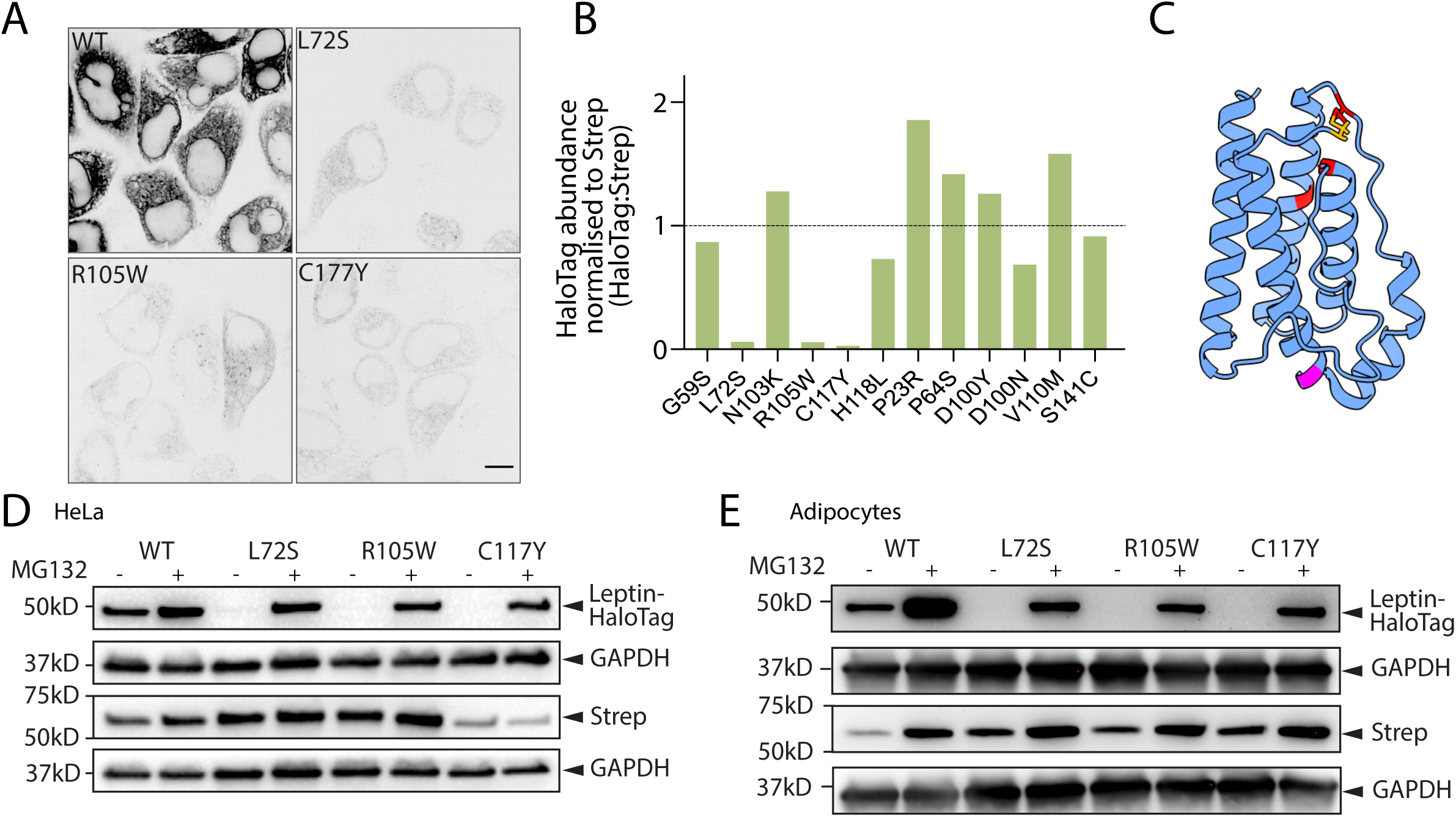
p.L72S, p.R105W, and p.C117Y leptin variations have low abundance due to proteasomal degradation. A) Fluorescence microscographs of HeLa cells expressing WT, p.L72S, p.R105W, and p.C117Y SBP-HaloTag-leptin before biotin addition. Representative of n= 7 for WT SBP-HaloTag-leptin, n = 3 for p.L72S, p.R105W, and p.C117Y SBP-HaloTag-leptin; scale bar: 10 µm. B) Quantification of leptin variant protein abundance normalised to Streptavidin-KDEL to account for plasmid integration efficiency and expressed relative to WT protein abundance (black dotted line). Data are n = 1. C) AlphaFold-generated model of leptin with signal peptide (amino acids 1-21) removed (39). Positions of variants that cause low leptin expression (L72S, R105W, C117Y) are in red. Position of S141C is in magenta. The position of the disulfide bond between C117 and C167 is in orange. D) HeLa cells expressing WT, p.L2S, p.R105W, or p.C117Y SBP-HaloTag-leptin were treated with 40 μM MG132 for 6 h where indicated, and lysates were analysed by Western blot. Streptavidin abundance was assessed as control for piggybac plasmid genomic insertion efficiency; GAPDH abundance was assessed as loading control. Representative blots of n= 3. E) 3T3-L1 adipocytes expressing WT, p.L2S, p.R105W, or p.C117Y SBP-HaloTag-leptin were treated with 40 uM MG132 for 6 h where indicated, and lysates were analysed by Western blot. Streptavidin abundance was assessed as control for piggybac plasmid genomic insertion efficiency; GAPDH abundance was assessed as a loading control. Representative blots of n = 3.

We reasoned that reduced protein abundance could arise from 1) poor genomic insertion efficiency during the stable cell line generation; 2) mRNA degradation; 3) decreased translation; 4) protein misfolding and subsequent degradation; or 5) aberrant sorting in the secretory pathway. To rule out low expression due to poor genomic integration, we assessed the abundance of both the SBP-HaloTag-leptin construct and Strep-KDEL. As the Strep-KDEL is on the same bicistronic vector as SBP-HaloTag-leptin, its expression reports on genomic integration and stability of the transgene. The SBP-HaloTag-leptin:Strep-KDEL ratio for p.L72S, p.R105W and p.C117Y was markedly reduced compared to WT SBP-HaloTag-leptin (**Fig. 4B**). Thus, our data support that p.L72S, p.R105W and p.C117Y variants are unstable at the mRNA or protein level (indicated in red in **Fig. 4C**).

Misfolded or unstable proteins retro-translocate from the ER using the ERAD pathway and are degraded in the cytosol by the proteasome (30). Therefore, we hypothesised that lower p.L72S, p.R105W and p.C117Y abundance was due to degradation. Consistent with this, treatment of cells with the proteasomal inhibitor MG132 increased the protein abundance of all three variants to approaching WT levels (**Fig. 4D**). Similarly, in cell lines generated in the more physiological 3T3-L1 adipocyte model, we observed the same low abundance of p.L72S, p.R105W and p.C117Y, which was also rescued by treatment with MG132 (**Fig. 4E**). Together, these data suggest that these variants are transcribed and translated normally, but are subsequently degraded by the proteasome resulting in low serum levels.

### p.S141C variant slows secretion due to improper ER retention

The p.S141C variant (indicated in magenta in **Fig. 4C**) was first identified in subjects with obesity from Turkmenistan (25), but there is currently no published data on serum leptin of carriers. p.S141C leptin had comparable initial fluorescence to WT leptin, suggesting that p.S141C leptin is produced and not degraded (**Fig. 3D**). However, the rate constant for leptin secretion was reduced by approximately half compared to WT (**Fig. 3D**). Fluorescence micrographs across the time series indicated that the p.S141C SBP-HaloTag-leptin was retained in the ER for longer (**Fig. 5A**) than WT leptin, consistent with slower secretion from the cell (**Fig. 3D**; **Supplemental Fig. 1**). We next validated the impaired secretory phenotype using a fluorescence plate assay to directly measure p.S141C leptin secretion into the culture medium. After 240 min, cellular p.S141C leptin content was 57% higher in cells expressing p.S141C than those expressing WT SBP-HaloTag-leptin, and media p.S141C SBP-HaloTag-leptin content was 25% lower (**Fig. 5B**). This corroborates data obtained from our screen (**Fig. 3C**; **Supplemental Fig. 1**) suggesting that p.S141C variant is secreted with slower kinetics than WT.

**Figure 5:**
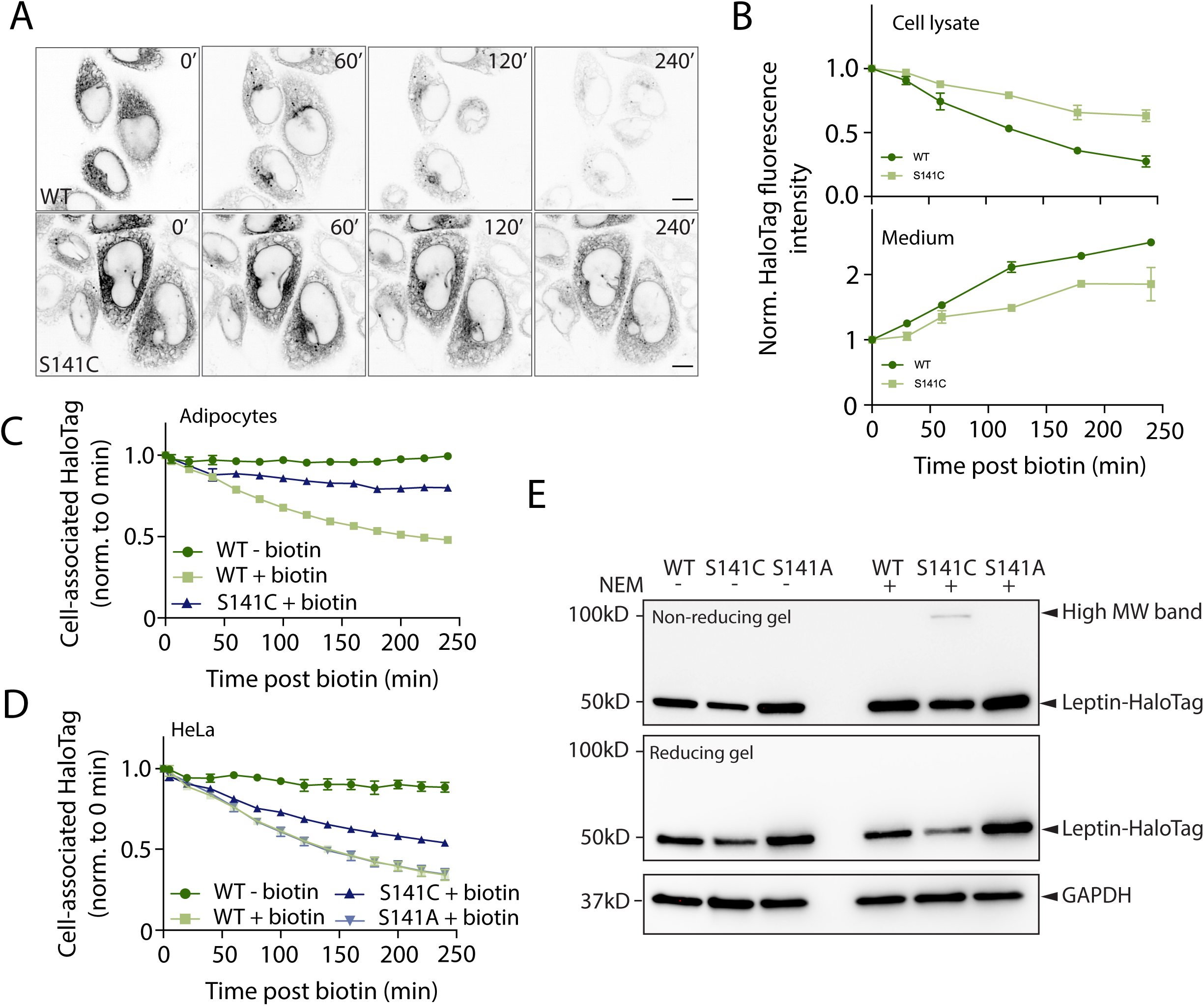
p.S141C variant is retained in the ER due to the additional Cys residue and forms a disulphide-linked complex. A) Fluorescence micrographs of HeLa cells expressing WT and p.S141C SBP-HaloTag-leptin before biotin addition, and after biotin at the indicated time points. Representative of n= 7 for WT SBP-HaloTag-leptin, n = 3 for p.S141C SBP-HaloTag-leptin. Scale bar: 10 µm. B) HeLa cells expressing WT or S141C SBP-HaloTag-leptin were treated with biotin for indicated times and the HaloTag fluorescence in cell lysate (top) and culture media (bottom) measured. n = 2. C) Kinetic secretory analysis of 3T3-L1 adipocytes expressing WT or p.S141C SBP-HaloTag-leptin treated with biotin where indicated. Cells were imaged at 20 min time intervals from 0-240 min. HaloTag fluorescence intensity was normalised to the 0 min time point. n = 3 for WT SBP-HaloTag-leptin; n = 2 for p.S141C SBP-HaloTag-leptin. D) Kinetic secretory analysis of HeLa cells expressing WT, p.S141C or p.S141A SBP-HaloTag-leptin treated with biotin where indicated. Cells were imaged at 20 min time intervals from 0-240 min. HaloTag fluorescence intensity was measured using an image analysis pipeline and normalised to 0 min time point. n = 2 for WT and p.S141C SBP-HaloTag-leptin; n = 3 for p.S141A SBP-HaloTag-leptin. E) Leptin variant molecular weight analysis by Western blot under reducing and non-reducing conditions. HeLa cells were treated with NEM to alkylate free cysteines and prevent disulfide bond formation or exchange after cell lysis. Anti-HaloTag antibody was used to detect SBP-HaloTag-leptin. A higher molecular weight anti-Halo-Tag antibody reactive band in cells expressing the p.S141C variant is indicated. GAPDH abundance was assessed as loading control. Data are representative of n = 4.

To test whether the p.S141C variant had altered secretory kinetics in a more physiological setting, we repeated the imaging-based leptin secretion assay in 3T3-L1 adipocytes. Again, the p.S141C variant exhibited slower secretion compared to WT (**Fig. 5C**).

We next asked if the loss of the serine or the gain of the cysteine was the cause of the secretory phenotype for the p.S141C leptin variant. We generated an additional p.S141A SBP-HaloTag-leptin variant. As previously (**Supplemental Fig. 1**), the p.S141C SBP-HaloTag-leptin variant exhibited slower secretion than WT SBP-HaloTag-leptin. However, the rate of secretion of p.S141A, as indicated by the decline in relative fluorescence intensity, was indistinguishable from WT leptin (**Fig. 5D**), suggesting that secretion was not impaired by the serine-to-alanine substitution. Therefore, slower secretion kinetics of the p.S141C is due to the gain of a cysteine residue.

WT leptin contains a disulfide between Cys117 and Cys167 (indicated in **Fig. 4C**) and a cysteine in the signal peptide at position seven that is cleaved during translocation (31). We hypothesised that the presence of an additional cysteine at 141 leads to the formation of an incorrect disulfide bond within leptin or with another protein. To test this indirectly, we treated cells with NEM to alkylate free cysteines to prevent disulfide bond formation or exchange after cell lysis and subjected cell lysates to non-reducing SDS-PAGE. Immunoblotting revealed the presence of a higher molecular weight anti-HaloTag immunoreactive band only in cells expressing the p.S141C variant (**Fig. 5E**). This higher molecular weight band was lost when lysates were reduced to break S-S bonds (**Fig. 5E**). These data suggest that the additional cysteine in p.S141C leptin-SBP-HaloTag causes the formation of a higher molecular weight disulfide-linked complex. Thus, our data are consistent with the addition of a cysteine in the p.S141C variant, resulting in aberrant disulfide bond formation, ER retention, and defective secretion due to protein quality control mechanisms in the ER.

## Discussion

Here, we established a novel cell-based assay to study the secretory kinetics of leptin. These studies in HeLa cells and adipocytes revealed that leptin is secreted via a classical route comprising the ER, Golgi, and post-Golgi vesicles, which fuse with the PM. Of the 12 leptin variants studies, three were post-translationally degraded by the proteasome (p.L72S, p.R105W, p.C117Y), and one was secreted at a slower rate than WT leptin (p.S141C). Further assays revealed that slower secretion of S141C was driven by ER retention as a result of the introduction of an additional cysteine and aberrant disulfide bond formation.

Understanding the regulation of leptin secretion can be challenging since constitutively secreted proteins are continually depleted from the cell under steady-state conditions. Additionally, 3T3-L1 adipocytes are widely used to study adipocyte biology but typically have low leptin mRNA levels (32, 33), making it difficult to study endogenous leptin secretion in these cells. Using the RUSH system, we have overcome these limitations by trapping ectopic leptin within the ER until its release *en masse* by the addition of biotin (17). By generating HeLa and cultured adipocyte lines, we have established the RUSH system in a line highly amenable to high-throughput assays and genetic manipulation and in a line that is more physiological.

Leptin was secreted via the classical ER-Golgi-PM secretory pathway in both HeLa cells and adipocytes. Further, the secretory kinetics of the SBP-HaloTag-leptin reporter used in this study were comparable to a control SBP-HaloTag construct. This suggests that leptin undergoes bulk secretion and does not contain signals to confer additional regulation on its rate of secretion. These data suggest that leptin secretion from adipocytes is most likely regulated transcriptionally (34) and translationally (9).

We selected 12 human leptin missense variants to study for effects on leptin expression and secretion. This included three variants that acted as negative controls, where the known effect of these variants is on leptin binding to its receptor (p.G59S, p.P64S, p.D100Y) and one where serum leptin is normal (p.V110M). We detected no effect of these variants on leptin protein abundance or secretion. However, four of the remaining nine variants were chosen for follow-up studies. One variant had slower secretory kinetics and longer ER retention compared to WT leptin. p.S141C (S120C in the mature protein after signal peptide cleavage) was identified in two subjects in Turkmenistan however, no leptin serum levels were reported (25). Our studies suggest that the appearance of the Cys residue causes increased ER retention and slower secretory kinetics, possibly through aberrant disulfide bond formation. Correct formation of disulphide bonds is a critical process for protein maturation and exit from the ER and is regulated by ER resident chaperones (35). WT leptin contains one disulphide bond between Cys117 and Cys167, which is necessary for secretion (31). Formation of this disulfide is critical for correct folding since the pC117Y variant, where this disulfide is disrupted, is degraded by the proteasome (**Fig. 4C**). Adiponectin, another secreted adipokine, is regulated by thiol-mediated retention by ERp44, an ER-localised chaperone that forms a mixed cysteine bond with adiponectin and retains it in the ER (36). ER-resident chaperones, such as ERp44, could form mixed disulfide bonds with the new free cysteine residue in the p.S141C variant, thereby retaining the protein in the ER. Alternatively, the free cysteine residue may lead to aberrant leptin dimer formation, as previously proposed for this variant (37), which may impede ER exit. Intriguingly, Hagluand et al., demonstrated that the p.S141C folds correctly and is stable and suggested that this leptin variant may interfere with forming an active quaternary complex with the leptin receptor (37). Our study provides an additional mechanism to p.S141C pathophysiology as we demonstrate it is defective in secretion. Therefore, we conclude that p.S141C alters systemic leptin responses through 1) decreased serum leptin (due to slow secretion) and 2) impaired receptor binding.

The other three leptin coding variants with low secretion resulted in proteasomal degradation of the leptin mutant protein (p.L72S, p.R105W, and p.C117Y). These data are entirely consistent with data from in vitro studies using these variants that showed these proteins, when purified from *E. coli,* were aggregated and/or misfolded (37). This suggests that, in the case of these variants, low serum leptin is driven by misfolding and post-translational degradation of leptin. We note that the conclusion contrasts with previous reports on the p.L72S and p.R105W variants, which have been reported to have increased cellular retention (12, 21) (i.e., impaired secretion). A possible reason for this disparity is that in these studies, HEK293T and Cos-1 cells were transiently transfected, which can result in excessive overexpression; if so, accumulation in the ER, prior to degradation, would be consistent with our data.

Of the six variants associated with low serum leptin (**Table 1**), we detected no change in either expression or secretory kinetics for two of them (p.N103K and p.P23R). For example, p.N103K was identified in a subject with obesity and very low serum leptin (22), yet expression and secretion were comparable to WT leptin in our studies. Possible explanations for these discrepancies between our data and clinical data include 1) subtle differences in expression of secretory kinetics that were not evident from our screen; 2) that these differences were masked by screening in a non-physiological cell type; 3) that coding variants of leptin artificially lower serum leptin detection through changes in ELISA antibody affinity; and 4) that the coding variants do not influence serum leptin and other genetic causes explain obesity in these cases.

Overall, in our study, we have developed cell models and a pipeline for studying the effect of *LEP* coding variants on leptin’s expression and secretory kinetics. From this, we were able to gain further insight into the mechanism of action *in vitro* for four documented variants of leptin, the post-translational control of leptin degradation, and its secretion. Our pipeline for studying secreted proteins using the RUSH system can be extended to other novel or existing leptin variants and is generalisable to other secreted proteins implicated in disease.

## Materials and Methods

### Molecular Cloning

pMPx92 (StrepKDEL, SBP-HaloTag) was generated as previously described (28). The human *LEP* mRNA sequence was sourced from the cDNA construct hLEP-pcDNA3.1(+)-C-(K)DYK (GenScript, clone ID OHu27387; NM_000230.3). Leptin-SBP-HaloTag plasmid was generated by inserting the WT *LEP* gene into the backbone derived from pMPx92 using restriction digest and Gibson assembly. The 13 mutant *LEP* sequences containing specific point mutations were generated from the original sequence by carrying out PCRs with mutagenic primers and using a KLD enzyme mix (M0554S; NEB) to repair the vectors. Sequences were confirmed using Sanger sequencing.

### Cell culture and transfection

HeLa cells were maintained in Dulbecco’s modified Eagle’s medium high glucose (DMEM; D6429; Merck supplemented with 10% foetal bovine serum (FBS; F7424; Merck) and containing MycoZap plus (VZA-2012; Lonza) at 37 °C and 5% CO_2_. 3T3-L1 cells grown in DMEM supplemented with 10% FBS and 1% GlutaMax (35050061; Gibco). at 37°C and 10% CO_2_. Stable HeLa cell lines were generated via transfection of HeLa cells with pMPx92-based vectors alongside pJEx21 containing the PiggyBac transposase, using Lipofectamine 2000 (11668019; Thermo Fisher), according to the manufacturer’s instructions. Stable 3T3-L1 cell lines were generated in a similar manner using the transfection reagent Viafect (E4981; Promega), according to the manufacturer’s instructions. Cell lines were selected using 250 µg/mL hygromycin (10843555001; Roche). 3T3-L1 fibroblasts were differentiated into mature adipocytes as previously described (38) and used in experiments 9-12 d after differentiation was initiated.

### Live cell confocal imaging

HeLa cells were plated onto Perkin Elmer 96-Well Phenoplate Ultra Plate (6055302; Perkin Elmer), 25x10^4^ cells per well, 24 h before imaging, in media containing JFX646 HaloTag-dye (200 nM; GA112A; Promega). When carrying out treatment with brefeldin A (BFA), media containing 10 μg/mL BFA was applied to the cells for 1 h before imaging. The DMEM was removed, the cells were washed with 1X phosphate-buffered saline (PBS), and the imaging media was applied. The imaging was carried out in FluoroBrite DMEM (A1896701; Gibco) containing 2% bovine serum albumin (BSA; A9418; Merck) and 1% GlutaMax. 3T3-L1 cells were similarly prepared, but the 96-well plate surface was coated with Matrigel (356234; Corning) prior to plating. Imaging was carried out on the Perkin Elmer Opera Phenix spinning disc confocal microscope equipped with a 63x/1.15 Water objective and environmental chamber (temperature and CO_2_ controlled at 37 °C and 5% CO_2_) with 2-pixel binning and 570-630 nm excitation and emission filters. A pre-treatment image was taken before the application of biotin-containing media to the cells. The cells were maintained at 37 °C and 5% CO_2_ and imaged every 20 minutes for 4 h, with an endpoint image taken 24 h after the application of biotin. Images were acquired using Harmony phenoLOGIC Software (v4.9; Perkin Elmer).

### Live cell super-resolution imaging

For live cell imaging, 9x10^4^ Leptin-SBP-HaloTag HeLa cells were plated onto matrigel-coated glass coverslips (CB00250RAC; Menzel-Gläser). After 48 h, cells were incubated for 1 h at 37 °C with fresh complete DMEM containing JFX646 HaloTag ligand (200 nM; GA112A; Promega), washed twice with PBS 1X and imaged on an Elyra 7 with Lattice SIM² microscope (Zeiss) equipped with an environmental chamber (temperature controlled at 37 °C, humidified 5% CO2 atmosphere), two PCO.edge sCMOS version 4.2 (CL HS) cameras (PCO), solid-state diode continuous wave lasers and a Zeiss Plan-Apochromat 63x/1.4 Oil DIC M27, all under the control of ZEN black software (Zeiss). D-biotin (B4501; Merck) at a final concentration of 500 µM was added to induce the RUSH.

For imaging of Leptin-SBP-HaloTag in adipocytes, cells were seeded onto matrigel-coated glass coverslips and imaged 12 d after differentiation. Imaging at the Elyra 7 with Lattice SIM² microscope was performed as described above.

### Quantification and analysis of microscopy data

The time series images acquired using the Opera Phenix were analysed using the Harmony phenoLOGIC Software (v4.9; Perkin Elmer). The total fluorescence was measured and averaged across six fields per well. Six wells were used for each cell line within a biological replicate, and three biological replicates were performed. The raw data from Harmony was processed in Microsoft Excel, where background fluorescence was subtracted, technical replicates averaged and normalised to their zero time points. Data was plotted, and statistical tests were performed using Prism GraphPad.

### Secretion of leptin in HeLa cells assay

Leptin–SBP-HaloTag HeLa cells were incubated with JFX646 HaloTag ligand (200 nM; GA112A; Promega) overnight. 1x10^6^ cells were aliquoted into 1.5 ml tubes and re-suspended in the DMEM supplemented with 10% FBS and 2.5% HEPES (H0887; Merck), and containing MycoZap plus. The cells were incubated at 37 °Cin a bench-top heat block, and D-biotin to a final concentration of 500 µM was added to induce the RUSH in a reverse time series, with regular vortexing of the cells to avoid settling. After stimulation, the cells were moved onto ice and centrifuged at 500 x g at 4 °C for 5 min. The supernatant was transferred to a 96-well plate (Perkin Elmer 96-Well Phenoplate Ultra Plate). The pellet was incubated with lysis buffer (10 mM Tris pH 8, 150 mM NaCl, 0.5 mM EDTA, 1% Triton) on ice for 20 min. The lysates were centrifuged for 10 min at 4 °C, 16,000 rpm, and the supernatant was transferred to the same 96-well plate. Fluorescence intensity was then measured using a Clariostar plus and analysed using Prism GraphPad.

### Western blotting

For analysis of proteasomal degradation, cells were treated with 40 μM MG132 (474790l Merck) for six h at 37 °C. Cells were washed twice with ice-cold PBS on ice and lysed in radioimmunoprecipitation assay (RIPA) buffer (10 mM Tris-HCl pH 8.0, 1 mM EDTA, 0.5 mM EGTA, 1% Triton X-100, 0.1% Sodium Deoxycholate, 0.1% SDS, 150 mM NaCl) containing proteinase inhibitors and sonicated. For N-Ethylmaleimide (NEM) (E3876; Merck) treatment, HeLa cells were washed twice in ice-cold PBS before incubation in cold PBS containing 100 mM NEM on ice for 10 min. Cells were then lysed in RIPA buffer containing 50 mM NEM and proteinase inhibitors and sonicated. Lysates were cleared by centrifugation for 10 min at 16,000 rpm at 4 °C, and protein concentration was determined using the BCA protein assay kit (A55864, Thermo Fisher Scientific). 10 µg of lysate was resolved via SDS-PAGE on a 4-20% gradient polyacrylamide gel (4561095; Bio-Rad) and transferred to a nitrocellulose membrane using the TransBlot Turbo mini-nitrocellulose kit (1704270; Bio-Rad). Membranes were blocked in 1X Tris-buffered saline containing 0.1% Tween-20 with 5% skimmed milk powder and immunoblotted as previously described. Antibodies used for blotting include anti-GADPH (clone 141C10, 2118S; Cell Signalling Technologies), anti-Streptavidin antibody (clone S10D4, MA1-20010; ThermoFisher Scientific), and anti-HaloTag antibody (G9221; Promega,) followed by incubation with horseradish peroxidase (HRP)-conjugated anti-rabbit or mouse immunoglobulin G (IgG).

## Acknowledgements

These studies were supported by the Institute of Metabolic Science Metabolic Research Laboratories Imaging Core (Wellcome Trust Major Award (208363/Z/17/Z)). The CIMR Flow Cytometry Core Facility and the CIMR Microscopy Facility supported this research.

## Author contributions

Conceptualisation (I.S.F., D.C.G., D.J.F.), methodology (D.C.G. and D.J.F.), formal analysis (H.J.M.B., A.S.S., D.S.), investigation (H.J.M.B., A.S.S., E.M.O., D.S., D.C.G., D.J.F., J.E.C., J.E.), writing of manuscript (H.J.M.B., A.S.S., J.E., D.C.G., D.J.F., with input from all authors), visualisation (H.J.M.B., A.S.S., D.C.G., D.J.F.), supervision (I.S.F., D.C.G., D.J.F.), project administration (I.S.F., D.C.G., D.J.F.).

## Rights Retention Statement

For the purpose of open access, the author has applied a Creative Commons Attribution (CC BY) licence to any Author Accepted Manuscript version arising.

**Supplementary figure 1. Leptin variant RUSH-secretion assay screen.** Kinetic secretory analysis of HeLa cells expressing WT or the indicated leptin variant. The same WT data is plotted on each graph for ease of comparison. Biotin was added where indicated. Cells were and imaged at 20 time intervals from 0-240 min. Cell-associated HaloTag fluorescence intensity was normalised to the 0 min time point. n= 7 for WT SBP-HaloTag-leptin; n = 3 for all leptin variants.

