## Supplementary figures and images for "A quantitative pipeline to assess secretion of human leptin coding variants reveals mechanisms underlying leptin deficiencies"

### Supplemental Figure 1

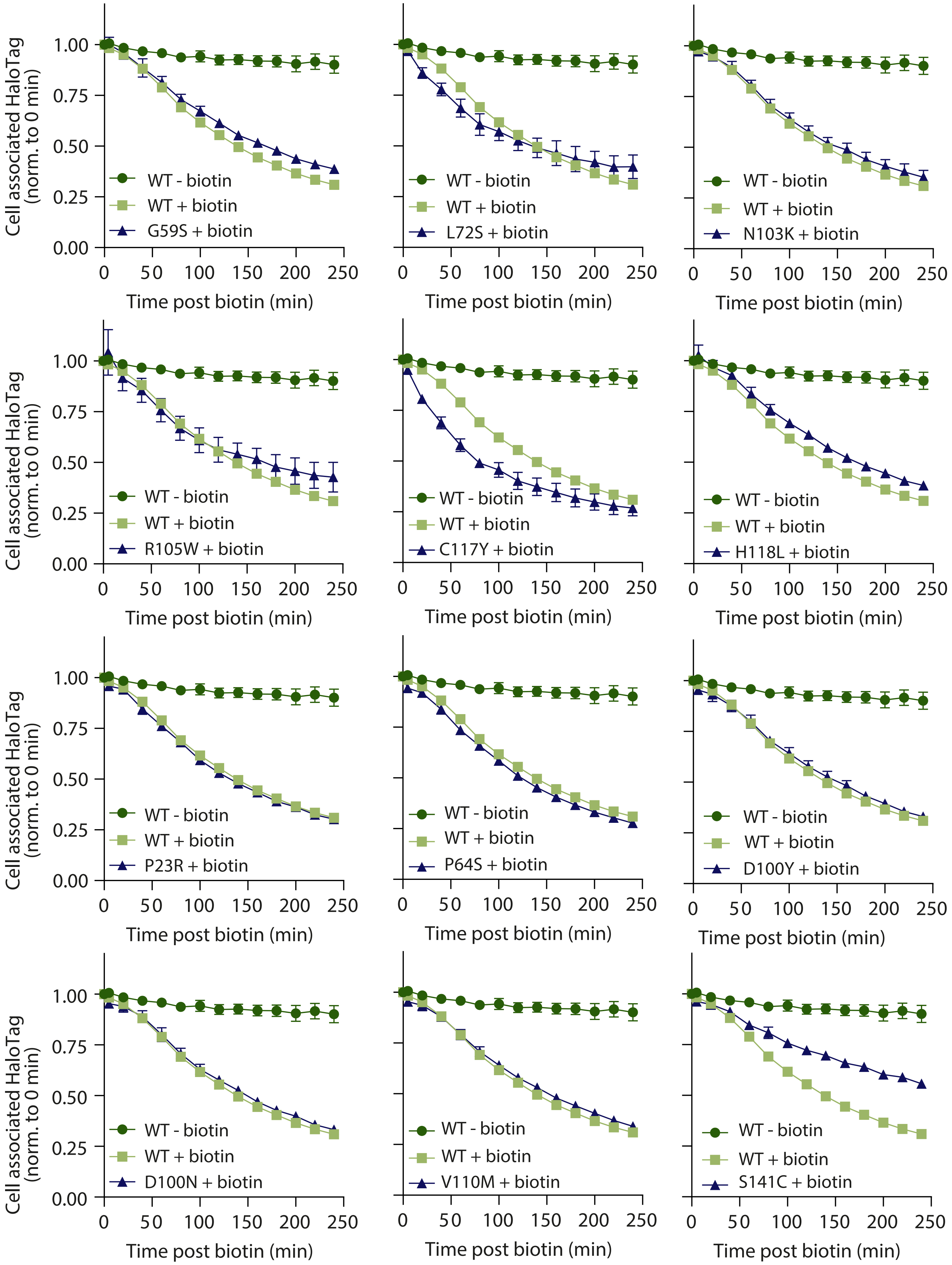
